# Distinct frontal cortical pathways for the long-range processing of auditory and visual stimuli

**DOI:** 10.64898/2026.08.22.746466

**Authors:** Elaida Dimwamwa, Amber M. Kline, Per-Niklas Barth, David M. Schneider

## Abstract

Frontal cortex neurons are sensory responsive and send long-range feedback to multiple different sensory cortices, but whether these functions are carried out by the same neurons or by distinct modality-specific circuits remains unknown. Using large-scale electrophysiology, two-photon calcium imaging, and viral circuit tracing in awake mice, we identify rich, modality-specific sensory coding in the frontal cortex (secondary motor/anterior cingulate cortex) that is largely dissociated from the neurons providing feedback to sensory cortex. Frontal cortex neurons exhibited robust sensory-evoked activity, with response magnitudes, latencies, and feature selectivity comparable to those observed in primary sensory cortex. Individual neurons displayed tuning for distinct sensory modalities, while population-level activity reliably decoded both sensory modality and stimulus identity. Anatomically, primary auditory (A1) and visual (V1) cortex axons were largely intermingled in anterior frontal cortex but more segregated in posterior regions, revealing spatial variation in the integration of sensory inputs. Frontal cortex neurons responsive to auditory stimuli were biased more anterior compared to visually-responsive neurons. Dual retrograde tracing identified distinct frontal cortex populations projecting back to A1 and V1 that were biased to the posterior and medial extent of the frontal cortex. A1-and V1-projecting frontal neurons were minimally sensory responsive, and no more likely to be responsive than other frontal neurons. Together, these findings reveal a division of labor within the frontal cortex, in which detailed sensory representations and corticocortical feedback arise from partially distinct neuronal populations. This circuit architecture provides a means through which specific sensory information can be transformed within the frontal cortex before being communicated back to the sensory cortex according to behavioral demands.

## Introduction

Flexible and context-specific behaviors are necessary for survival, requiring the brain to integrate distinct sensory information about the external world with internal and behavioral state. Top-down projections from frontal cortical regions including the secondary motor cortex (M2) and the anterior cingulate cortex (ACC) provide an important route by which such internal and behavioral variables can modulate the bottom-up processing of sensory input in primary sensory regions (Crapse and Sommer, 2008; Niell and Stryker, 2010; Keller et al., 2012; Schneider et al., 2014, 2018; Cullen, 2023). In addition to sending behavior-related signals to sensory cortex, neurons within the frontal cortex are also sensory responsive, suggesting that sensory information is available within circuits traditionally associated with behavioral and motor control (Schall, 2015; Holey and Schneider, 2024; Lien and Haider, 2025).

The reciprocal organization of these connections raises an important question about how distinct sensory information is incorporated into frontal circuits and subsequently routed back to the sensory cortex. The frontal cortex could provide a common behavioral-related signal to multiple sensory systems, effectively broadcasting information about the animal’s state across sensory cortices. Alternatively, distinct frontal populations could route modality-specific information to individual sensory systems. In support of the latter possibility, frontal activity can be selectively shaped by specific sensory experiences. In the visual system, altering the relationship between self-motion and visual feedback changes activity in ACC projections to primary visual cortex (V1) and contributes to mismatch signaling in V1, even when the animal’s motor behavior remains similar (Leinweber et al., 2017). Likewise, M2 neurons change their responses when movements produce expected versus unexpected auditory feedback (Holey and Schneider, 2024). These findings suggest that frontal feedback can incorporate specific, sensory-related information rather than simply reflecting a generic movement signal, raising the possibility that frontal-sensory interactions are organized into distinct pathways for different sensory modalities.

The anatomical and functional organization of the reciprocal frontal-sensory interactions for distinct sensory modalities remains unknown, which we address here using complementary anatomical tracing, electrophysiology and two-photon calcium imaging. We found that auditory and visual information is robustly represented within the frontal cortex, and that long-range connections from the primary auditory (A1) and visual (V1) cortices exhibit distinct anatomical and spatial organization. We also find that the populations of frontal cortical projectors to A1 and V1 inputs largely do not overlap nor are they preferentially sensory responsive. Together these findings reveal an organization of frontal-sensory circuits that combines broadly distributed multimodal sensory representations with anatomically distinct pathways for communication with individual sensory cortices.

## Results

### Frontal cortex neurons distinctly encode auditory and visual input

To determine the extent to which frontal cortex neurons respond to auditory versus visual input, we performed multi-channel silicon probe recordings of populations of frontal cortical neurons in awake, head-restrained mice passively receiving auditory and visual input (see Methods, Figure 1a). We targeted our recordings to both the secondary motor cortex (M2) and the anterior cingulate cortex (ACC) because each of these areas have been individually previously established as frontal nodes for reciprocal interactions with the auditory and visual cortex, respectively (Nelson et al., 2013; Schneider et al., 2014, 2018; Leinweber et al., 2017; Zhou and Schneider, 2024; Lien and Haider, 2025). We first presented an auditory noise stimulus and a visual flash stimulus, designed to span a broad range of feature space for each modality. Consistent with previous reports, we observed robust sensory responses to both the auditory and visual stimuli (25% and 15% of neurons respectively, for auditory and visual stimuli; Figures 1b-c). Most neurons responded by increasing their firing rate (22% and 14% for the noise and flash, respectively), while a smaller subset responded by decreasing their firing rate (3% and 1% for the noise and flash, respectively).

**Figure 1.**
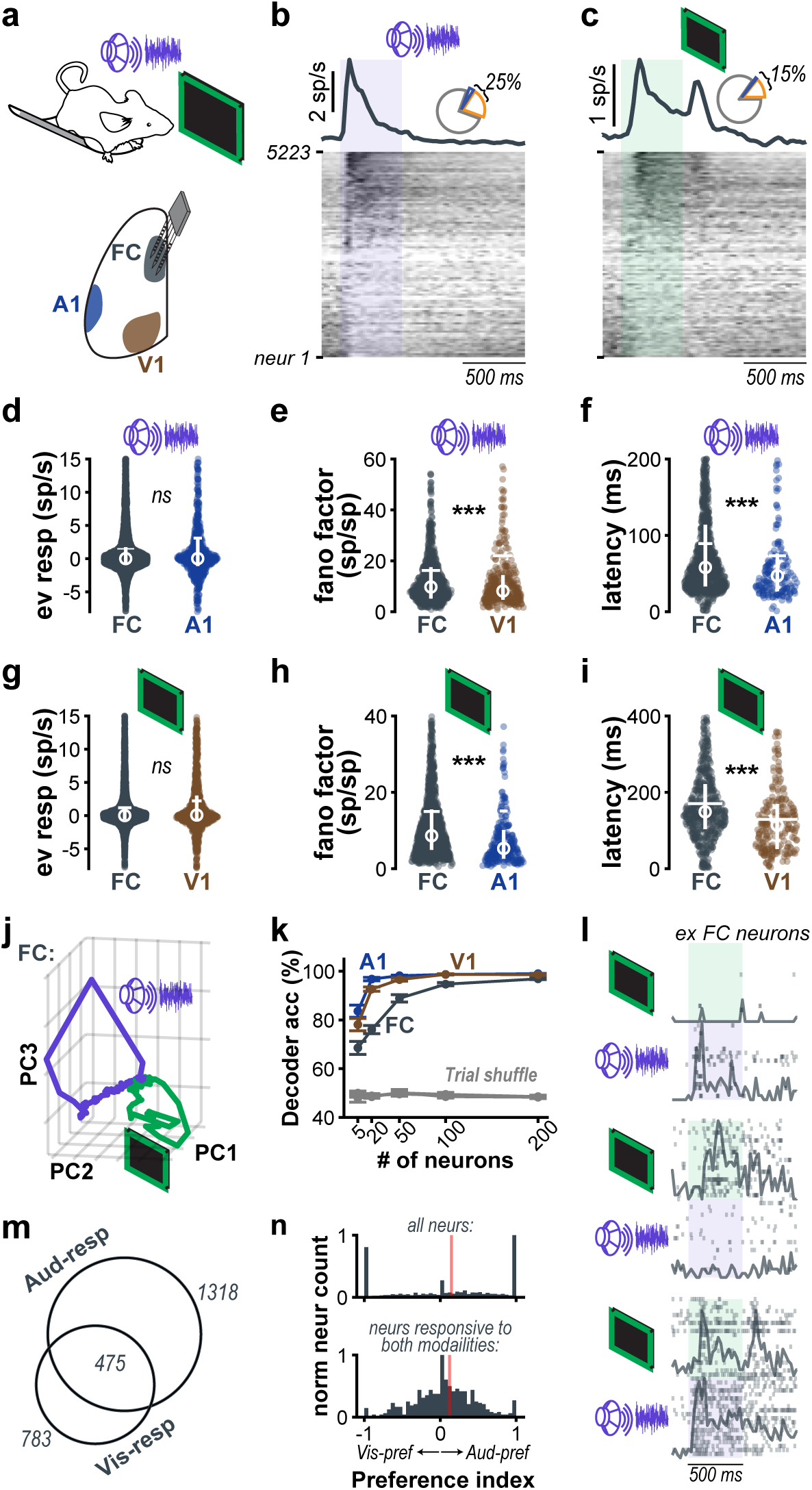
Frontal cortical neurons distinctly encode auditory and visual input. **(a)** We performed multi-channel, silicon probe recordings targeted to the frontal cortex (FC), the primary auditory cortex (A1), and the primary visual cortex (V1) in awake and head-fixed mice receiving auditory and visual input. **(b)** Frontal cortex population peri-stimulus time histogram (psth, top) and individual neuron psths (bottom, normalized to peak of each neuron) to an auditory noise stimulus. The shading indicates the stimulation period. Pie chart inset: Proportion of FC neurons that responded to the auditory noise stimulus with an increase in spiking (orange), a decrease in spiking (blue), or did not respond (gray). **(c)** Same as ***b***, but in response to a visual flash. **(d)** Auditory noise-evoked response of FC and A1 neurons (p = 0.89). The horizontal line indicates mean across neurons, the circle indicates the median, and the vertical line indicates the first to the third interquartile range (n = 5223 FC neurons, 661 A1 neurons, 1170 V1 neurons). **(e)** Fano factor of auditory noise-responsive A1 and FC neurons (p = 1.5e-18). **(f)** Response onset latency of auditory noise-responsive A1 and FC (p = 2e-6). **(g-i)** Same as ***d-f***, but for FC and V1 in response to the visual flash (evoked response: p = 0.12; fano factor: p = 6.9e-12; latency: p = 7.6e-9). **(j)** Principal component analysis of the trajectories of frontal cortex neurons in response ot the auditory noise (purple) and visual flash (green). **(k)** Mean +/-sem support vector machine decoder accuracy using the actiivity of subsets of neurons of varying size from each cortical region. **(l)** Example responses of simultaneously recorded frontal cortical neurons with differing selectivity to the auditory noise vs the visual flash. The shaded box indicates the period of stimulation. **(m)** The number of frontal cortical neurons that are responsive to only the auditory flash, only the visual flash, or both. **(n)** Normalized histograms of the preference index of frontal cortical neurons. The red line indicates the mean of each histogram.

To benchmark the auditory and visual responses in frontal cortical neurons, we also recorded from populations of neurons in the primary auditory cortex (A1) and the primary visual cortex (V1) of awake, head-restrained mice under the same conditions. Intriguingly, the baseline-subtracted, stimulus evoked response to the auditory noise stimulus of individual frontal cortical neurons was statistically indistinguishable from the responses in A1 (Figure 1d). However, the frontal cortical responses to auditory noise were generally more variable across trials than those in A1 (Figure 1e). Although frontal cortical neurons were on average slower to respond than A1 neurons by 11 ms, some frontal cortical neurons had onset latencies that were as short as those observed in A1 (Figure 1f). Similarly, the visual responses across the entire population of frontal cortical neurons were statistically indistinguishable and more variable than those in V1 (Figures 1 g-h). Here again, we found that while the responsive frontal cortical neurons responded, on average, 41 ms slower than V1 (Figure 1i), many neurons responded with comparably fast latencies to those in V1, corroborating previous reports (Lien and Haider, 2025).

We next sought to determine whether the auditory and visual stimuli were distinctly represented in the frontal cortex. We first examined the population dynamics evoked by each modality by applying principal component analysis (PCA) on the frontal cortical activity. We visualized the population trajectory in the first three principal components (which account for 16.5%, 7.0%, and 4.4% of the variance explained, respectively) and observed distinct frontal cortical population trajectories in response to each stimulus modality (Figure 1j). This suggests that the encoding of auditory and visual inputs is separable at the population-level. To further verify this, we built a support vector machine (SVM) to decode the modality of each stimulus from stimulus-evoked spike counts of subsets of neurons in the population. We found that even with as few as five frontal cortical neurons, we could decode the modality of the stimulus above chance, with decoding accuracy plateauing using around 100 neurons. The same decoding approach applied to A1 and V1 neurons outperforms the decoding on frontal cortical neurons using smaller pools of neurons, distinguishing frontal cortical sensory encoding from that of primary sensory regions (Figure 1k).

At the single cell level, simultaneously recorded neurons showed varying degrees of selectivity for auditory and visual stimuli (Figure 1l). We observed this mixing of the preference of individual frontal cortical neurons across all recording sites such that across the population of recorded neurons, there were 1,1113 neurons that were responsive to the auditory noise, 546 that were responsive to the visual flash, and 316 that were responsive to both stimuli (Figure 1m). We measured the preference index of each frontal cortical neuron and found a distribution with strong tails, confirming that many frontal cortical neurons respond to only 1 stimulus modality, with an overall preference for the auditory stimulus. Even among the frontal cortical neurons that respond to both stimuli, the population was skewed towards preferring auditory stimuli (Figure 1l). To verify that this apparent modality selectivity in frontal cortical neurons was not simply a result of stimulus selection, we presented other simple and complex auditory and visual stimuli. While the stimulus choice does change the set of neurons that are responsive, many neurons continue to respond to stimuli of a single modality (Figure S1).

### Stimulus-evoked responses in frontal cortex cannot be fully explained by movement

It has previously been suggested that auditory responses in a non-auditory region such as V1 are more likely related to stimulus-evoked movement, rather than being stimulus-evoked, *per se* (Bimbard et al., 2023). In contrast, other recent work has shown that auditory and visual responses in the frontal cortex are reduced when the auditory or visual cortex are suppressed, suggesting that stimulus-evoked activity in the frontal cortex cannot be accounted for by movement alone (Holey and Schneider, 2024; Lien and Haider, 2025). To determine whether frontal cortex sensory responses can be accounted for by movement, we performed simultaneous videography of the mouse’s face and body during stimulus presentation (Figure 2a). We focused on a region of the face caudal to the vibrissal array that reliably encodes stereotyped, sound-evoked twitches (Figure 2b; Clayton et al., 2024).

**Figure 2.**
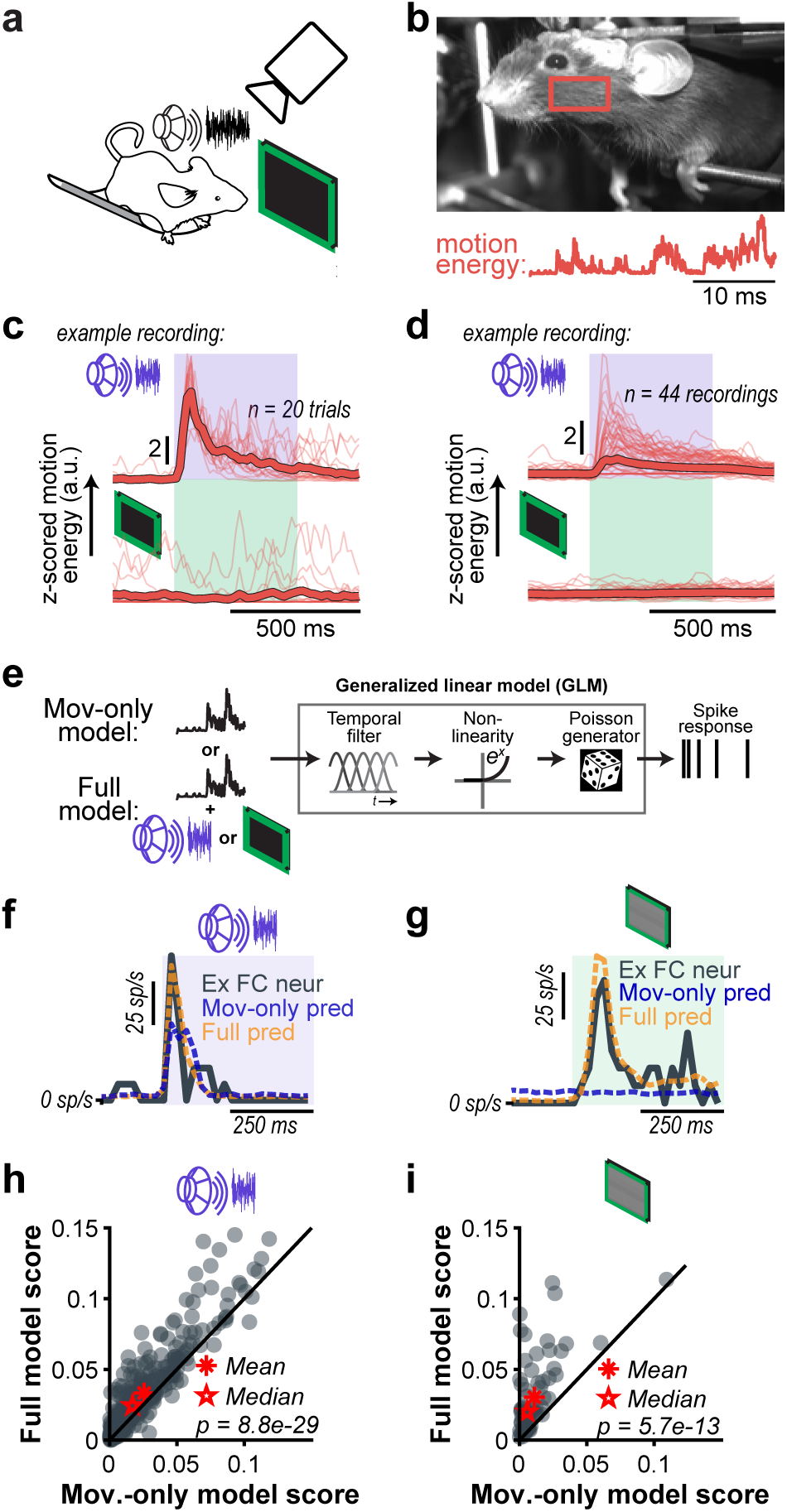
Stimulus-evoked responses in the frontal cortex cannot be fully explained by movement. **(a)** We used an infrared camera to monitor the neuronal activity in the same head-restrained and awake mice that were passively hearing auditory noise and viewing visual flashes in Figure 1. **(b)** To assess stimulus-evoked movement of each mouse during stimulus presentation, we extracted the motion energy of the outlined region of interest that is caudal to the vibrissa array. **(c)** The auditory noise-and visual flash-evoked motion energy of an example recording. **(d)** Same as **c**, but for all recordings. **(e)** We modeled the sensory stimulus-evoked response of individual neurons using a generalized linear model (GLM). Two models were built for each neuron, a movement-only model that only to the motion energy as input predictors of the neuron’s spiking, and full model that took both motion energy and the stimulus as predictors. **(f)** The auditory noise-evoked response of an example frontal cortical neuron overlaid with the predictions of the two models. **(g)** Same as **f**, but for an example neuron in response to the visual flash. **(h)** The McFadden’s pseudo-R^2^ scores to evaluate the performance of each model fit to predict the response to the auditory noise of responsive neurons only. Poorly fit neurons with a score less than 0 were excluded from the analysis (n=341). **(i).** Same as h, but for models fitting the response to the visual flash (n=69).

From this ROI, we find that auditory noise, but not the visual flash, reliably increases the motion energy across trials (Figures 2c-d). These data suggest that some auditory responses may be due to movement rather than sensation. To more rigorously address this, we next sought to determine the extent to which movement explains our observed responses by fitting generalized linear models (GLM) to the activity of single neurons (Nelder and Wedderburn, 1972; Pillow et al., 2005). For each neuron, we fit two models, one that accounted for movement as well as stimulus onset (i.e. “full model”) and one that only considered movement (“movement-only model”; Figure 2e). We assessed the extent to which the full model better predicted the sensory response of each sensory-responsive neuron compared to the movement-only model. Figures 2f and 2g show the response of frontal cortical neurons that are responsive to the auditory noise and visual flash, respectively, overlaid with the prediction of each model. The predictions for both neurons were substantially better for the full model compared to the movement-only model. We measured the improvement in model fit for the full model vs the movement-only model by comparing McFadden’s pseudo-R^2^ score, with larger numbers indicating improved fit. While the activity of some neurons was well accounted for by movement alone, across the entire population of responsive neurons, the full model performance was significantly better than the movement-only model performance (Figures 2h-i), consistent with the observed stimulus-evoked responses in the frontal cortex reflecting true sensory responses.

### Frontal cortex encodes specific acoustic and visual features

As further verification that the responses we measured contain a sensory component, we next presented pure tones of varying frequency and drifting gratings of varying orientation. Pure tones drove less stereotyped and smaller amplitude facial motion energy than the auditory noise, consistent with previous reports (Clayton et al., 2024; Figures 3a-b). Drifting gratings also do not drive stereotyped increases in motion energy. Nonetheless, we observed robust responses to both the auditory pure tones and visual gratings in the recorded frontal cortical population (Figures 3c-d). Notably, simultaneously recorded frontal neurons, which experienced the same stimulus-evoked movements, exhibited markedly different tuning across tones and grating orientations (Figures 3e–f). The presence of neuron-specific feature selectivity under a shared motor context argues against movement as the sole source of these responses and supports genuine sensory encoding in the frontal cortex.

**Figure 3.**
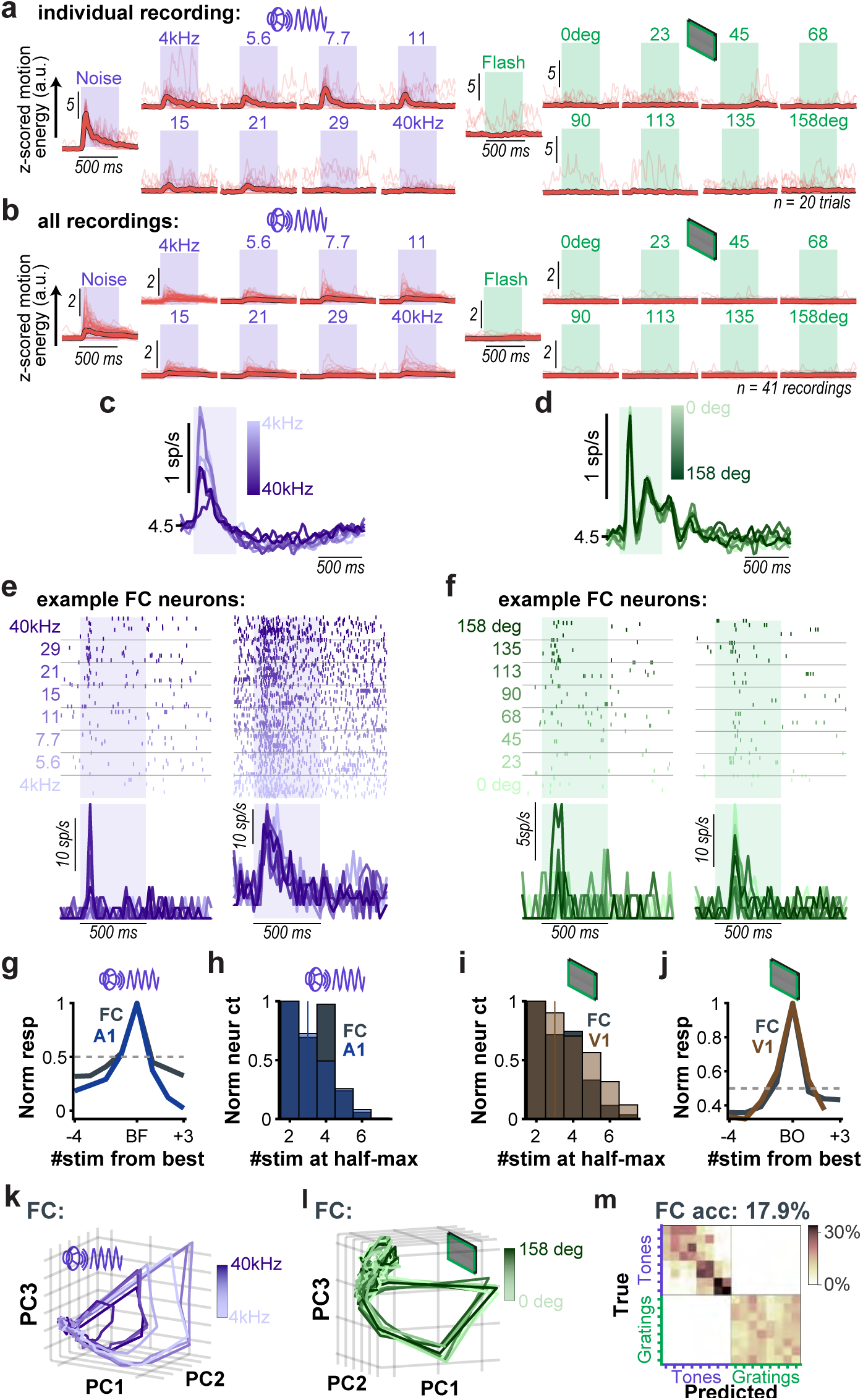
Frontal cortex encodes specific acoustic and visual features. **(a)** Stimulus-evoked motion energy for all stimuli for an example recording (thin lines: individual trials; thick lines: mean; horizontal and vertical scales for all plots are the same). **(b)** Same as **a**, but for all recordings (thin lines: mean of individual recordings; thick lines: mean across recordings). **(c)** Frontal cortex population PSTHs to one of eight auditory pure tones. **(d)** Same as **c**, but to one of eight visual drifting gratings. **(e)** The response of two simultaneously recorded frontal cortical neurons with different tuning to the auditory pure tones. **(f)** Same as e, but for the visual gratings. **(g)** The average tuning curve for the cortical populations. **(h)** Histogram of the number of stim at half-max from the tuning curve of individual neurons from each cortical population. **(i-j)** Same as **g-h**, but for the visual grating. **(k)** The frontal cortical population trajectory for each of the eight presented pure tones. Dimensionality reduction was performed using principal component analysis (PC1: 8.8% variance explained (ve); PC2: 4.2% ve; PC3: 1.9% ve). **(l)** Same as k, but for the visual gratings (PC1: 5.7% ve; PC2: 1.9% ve; PC3: 1.1% ve). **(m)** SVM decoding of the 16 stimuli from the stimulus-evoked, trial-by-trial spike counts of individual frontal cortical neurons (Chance: 6.25%).

We benchmarked the tuning of all recorded frontal neurons to that of A1 and V1 neurons recorded under the same conditions and found that the frontal cortical population has comparable tuning to pure tones as A1 neurons and comparable tuning to drifting gratings as V1 neurons (Figures 3g-j). PCA analysis applied to the frontal population unveils separation of the population trajectory for individual tones and gratings, respectively (Figures 3k-l). A decoder tasked to identify 1 of the 16 stimuli (8 auditory, 8 visual) from the frontal population does so significantly above chance levels and, importantly, the decoder never confuses an auditory stimulus for a visual one (Figure 3m).

Collectively, these data show that frontal cortex neurons distinctly encode multimodal sensory information and have sensory response properties on par with those observed in primary sensory cortex, including short response latencies, feature-specific receptive fields, and population representations that resolve distinct stimuli.

### Auditory and visual anatomical inputs to frontal cortex are spatially non-uniform

We next asked whether the functional segregation of auditory and visual responses in the frontal cortex is supported by distinct long-range input pathways from sensory regions. To test this, we mapped the distributions of axons arising from A1 and V1 that terminate in the frontal cortex. We injected adeno-associated viruses (AAVs) to express either EGFP or mCherry in A1 or V1 neurons, respectively (Figures 4a-b). We then imaged the A1 and V1 terminal fields in coronal sections throughout a broad anteroposterior range of the frontal cortex (-0.5 to +2.2 mm relative to bregma; Figure 4c). These anatomical analyses allowed us to examine a broader range of frontal cortex than was sampled using electrophysiology, including more lateral frontal cortex (M1) and a broader swath of ACC.

**Figure 4.**
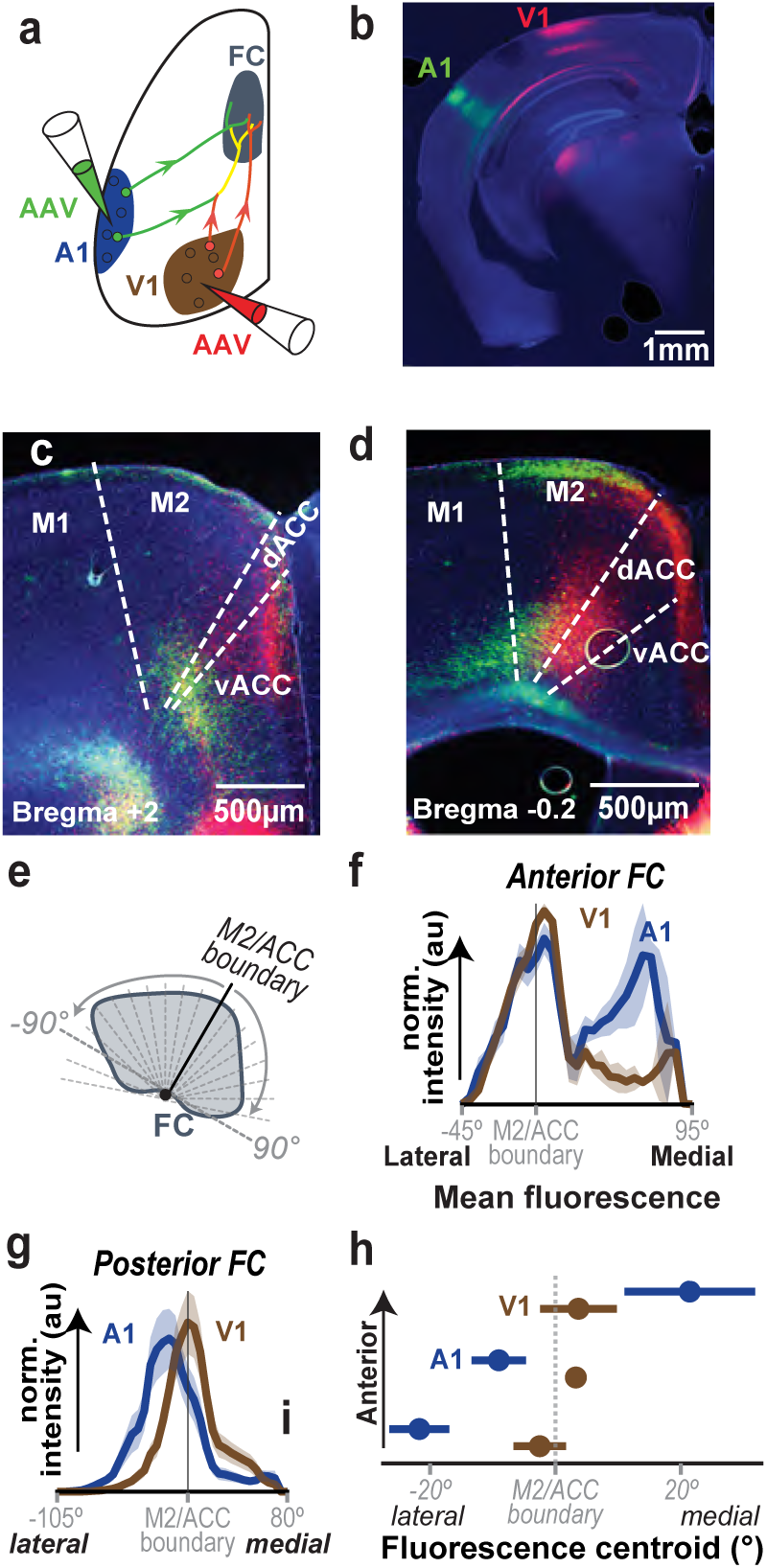
Auditory and visual anatomical inputs to frontal cortex are sptially non-uniform. **(a)** Dual anterograde labeling of A1 and V1 terminal fields in frontal cortex. **(b)** Coronal slice showing the A1 (green) and V1 (red) injection sites. **(c)** Example brain slice from an anterior section of frontal cortex, +2 Bregma. Cortical sub-region were demarcated by overlaying the appropriate atlas slice. **(d)** Same as ***(c)*** for a posterior slice,-0.2 Bregma. **(e)** Schematic demonstrating how the spatial analysis was conducted. For each slice, the M2/ACC boundary was defined, and the rest of frontal cortex was divided into 5° wedges. The intensity of each wedge was normalized to the wedge with the maximum intensity, and then intensity across slices was averaged. **(f)** Mean +/-SEM intensity distribution of A1 and V1 terminal fields in anterior regions of frontal cortex (N = 3 mice, n = 8 slices). **(g)** Same as ***(f)*** for posterior regions of frontal cortex. **(h)** Weighted median centroid per slice +/-standard deviation of A1 and V1 axon distributions in anterior, middle, and posterior slices of FC.

We found labelling of A1 and V1 axons throughout the entire antero-posterior range of the frontal cortex, with many individual slices showing a spatial segregation of A1 and V1 axons (Figure 4c-d). To quantify the spatial separation, we measured the density of A1 and V1 axons across the mediolateral axis. To account for the curvature of the cortex close to the midline, we divided the cortex into 5° radial wedges originating at the boundary of M2 and ACC and quantified the fluorophore density within each wedge (Figure 4e). Axons originating from V1 were most dense along the boundary between M2 and ACC, and this mediolateral organization was maintained along the anteroposterior range of the frontal cortex. In contrast, while A1 axons largely overlap with V1 axons in anterior frontal cortex, their distribution shifts medially in more anterior slices of the frontal cortex (Figures 4f-h).

These anatomical data reveal that auditory and visual inputs to the frontal cortex are promiscuous but are not homogeneously distributed throughout the entirety of the frontal cortex. Instead, A1 and V1 axons are more intermingled in anterior regions, while they become more separated in posterior regions. This anatomical distribution could support the combination of mono-and multimodal sensory processing observed during physiology.

### Auditory-and visual-encoding neurons in the frontal cortex are locally intermingled, but globally organized

Given that auditory and visual inputs overlap in anterior frontal cortex but begin to segregate in posterior regions, we next asked whether neuronal responses to different modalities displayed similar spatial organization. We used two-photon calcium imaging in awake, head-restrained mice to measure the sensory responses of frontal cortex neurons expressing GCaMP8s (Figure 5a). We imaged neurons in layer 2/3 in neighboring fields-of-view that were subsequently stitched together, yielding a composite imaging area spanning approximately 1.2 mm in the anteroposterior axis and 800 µm in the mediolateral axis (Figure 5d, see Methods). Across this large cortical region, consistent with the results of our electrophysiological recordings, we observed strong neuronal responses to both auditory and visual cues (Figure 5b). Individual neurons were frequently selective for only one modality, and within an imaging session, distinct neurons were responsive to stimuli with distinct features (Data not shown). The fraction of sensory-responsive neurons and the ratio of auditory and visual responsive cells were similar to those observed using electrophysiology (Figure 5c; auditory-responsive = 19.0%, visual-responsive = 13.7%).

**Figure 5.**
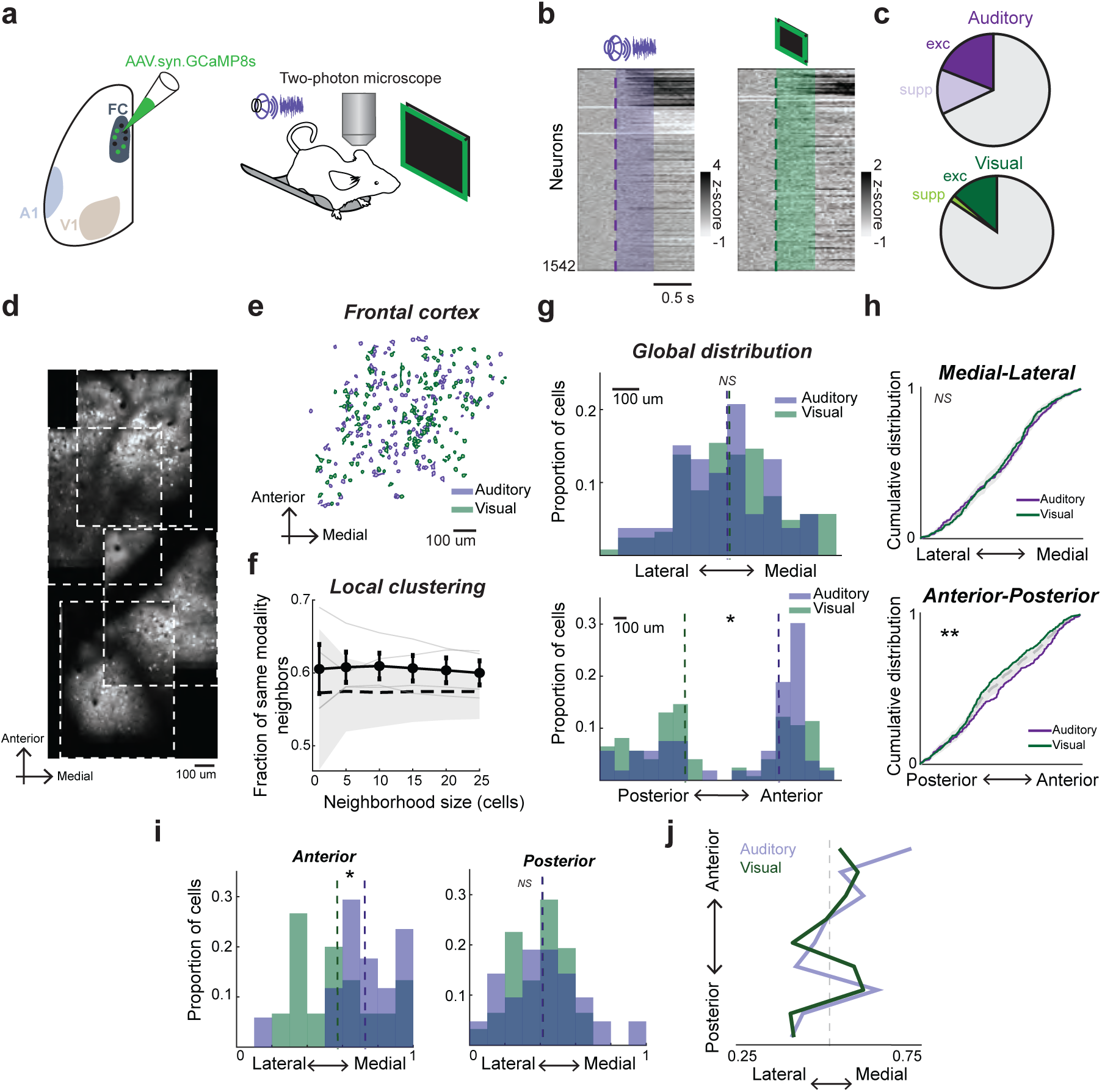
Auditory-and visual-responsive neurons display a spatial gradient in frontal cortex. **(a)** Left, AAV9.syn.GCaMP8s was injected into left frontal cortex. Right, two-photon calcium imaging was performed in awake, head-fixed mice while they were presented with randomized trials of pure tones, white noise, flashes and drifting gratings. **(b)** Sensory responsive neurons in example mouse across multiple fields-of-view in frontal cortex. n = 1542 neurons. **(c)** top, fraction of neurons excited (n = 293 cells, 19.0%) and suppressed (n = 201 cells, 13.%) by auditory cues; bottom, fraction of neurons excited (n = 212 cells, 13.7%) and suppressed (n = 28 cells, 1.8%) by visual cues. **(d)** Example stitching of frontal cortex fields-of-view. **(e)** Example cell masks of auditory-(purple) and visually-responsive (green) cells in frontal cortex. **(f)** Quantification of average fraction of same-modality nearest neighbors as a function of neighborhood size (k = 1-25). Solid black line represents the group observed mean +/-SEM across mice (n = 4 mice). Individual mice are plotted in gray. The gray shaded region indicates the 95% confidence interval of the null distribution generated via spatial label perumtation (n = 500 shuffles). p > 0.05 for all neighborhood sizes, Wilcoxon signed-rank test. **(g)** Distribution of sensory responsive frontal cortex neurons divided by modality across medial-lateral (ML, top) and anterior-posterior (AP, bottom) space in example mouse. Dashed lines represnt group medians. ML: auditory median = 337 um from midline, visual median = 330 um from midline, p = 0.8. AP: auditory median = 376 um from anterior edge, visual median = 819 um from anterior edge, p = 0.018, Wilcoxon signed-rank test. **(h)** Cumulative fraction of auditory (purple) and visual (green) neurons as a function of patial coordinate along the ML (top) and AP (bottom) axis (ML: p = 0.76; AP: p = 0.0048, two-sample Kolmogorov-Smirnov test. Gray dashed line and shading represent 95% confidence interval from shuffling spatial labels. **(i)** Proportion of auditory (purple) and visual (green) neurons along the normalized ML axis, pooled across mice (n = 4), divided by the most anterior (left) and posterior (right) quarter of frontal cortex space. Dashed lines represent group medians. Anterior: auditory median = 0.73, visual = 0.56, p = 0.039; Posterior: auditory median = 0.41, visual = 0.41, p = 0.823, Wilcoxon signed-rank test. Distributions of auditory and visual populations are significantly different in the anterior section but not the posterior region (anterior: p = 0.008; posterior: p = 0.815, two-sample Kolmogorov-Smirnov test). Both auditory and visual ML distributions are significantly different in anterior and posterior regions of frontal cortex (aud: p < 0.0001; vis: p = 0.028, two-sample Kolmogorov-Smirnov test). **(j)** Continuous average normalized ML position plotted as a function of the AP axis for auditory (purple) and visual (green) populations across animals. Lines represent median of each population across AP bins. n = 4 mice.

We next sought to determine the spatial organization of sensory-selective neurons in the frontal cortex, both at the local and global level. To measure fine-scale spatial organization, we evaluated if sensory-selective neurons are more likely to be near other neurons that prefer the same modality. Regardless of whether we looked at few or several neighbors (k = 1-25), we found that auditory and visually-responsive neurons were highly intermingled and that the observed fraction of same modality nearest neighbors were similar to the null distribution from shuffled modality labels (Figures 5e-f, p>0.05). Despite this “salt-and-pepper” organization at the local level, we observed a clear global spatial bias, with auditory-responsive neurons shifted toward more anterior regions of the frontal cortex relative to visually-responsive neurons (Figures 5g-h). Consistent with anatomical tracing experiments, we found that the medial-lateral distribution of auditory-and visually-responsive neurons depended on the anteroposterior position of the FOV, such that sound-responsive neurons are biased medially in anterior regions of the field of view, but laterally in posterior regions (Figure 5i-j). Thus, sensory representations in the frontal cortex exhibit a hierarchical spatial organization in which neurons encoding different modalities are locally intermingled but show broader spatial biases across the cortical landscape. Notably, despite differences in spatial scale, this spatial organization of sensory responses aligns with the anatomical organization of auditory and visual inputs to the frontal cortex.

### A1-and V1-projecting frontal cortex neurons are distinct populations, but share spatial biases

In addition to receiving inputs from sensory regions, the frontal cortex also provides top-down feedback to both A1 and V1, forming reciprocal loops (Nelson et al., 2013; Zhang et al., 2014). Given our functional and anatomical findings of modality-specific routing, there are several possibilities for how top-down feedback to different sensory cortices might be organized. Two features must be considered: first, do output neurons project exclusively to one sensory cortex, or broadcast signals to multiple modalities? Second, what type of information do output neurons encode? A hierarchical view of sensory processing predicts that frontal neurons projecting to a given sensory cortex would preferentially encode information from that same modality. Alternatively, these neurons could integrate information across modalities, allowing top-down feedback to reflect a multimodal sensory representation. A third possibility is that sensory-projecting frontal neurons encode little sensory information themselves, instead conveying nonsensory variables related to the internal or behavioral state, which could be relevant or irrelevant to the sensory experience.

To begin to distinguish between these models and determine whether top-down feedback to the sensory cortex is also modality-specific, we first assessed the specificity of the sensory targets. We injected retrograde AAVs into A1 and V1 to express either EGFP or mCherry in frontal cortical neurons that project to each region, randomizing the fluorophore region pairing per mouse to control for virus affinity (Figure 6a). We then confirmed expression of each fluorophore in the injection sites in A1 and V1 before imaging the cell bodies in the frontal cortex (Figures 6b-d). Interestingly, we observed largely non-overlapping populations of A1 and V1 projectors, with fewer than 3% of labeled cells projecting to both areas (Figure 6e), providing strong support for distinct frontal feedback pathways across modalities.

**Figure 6.**
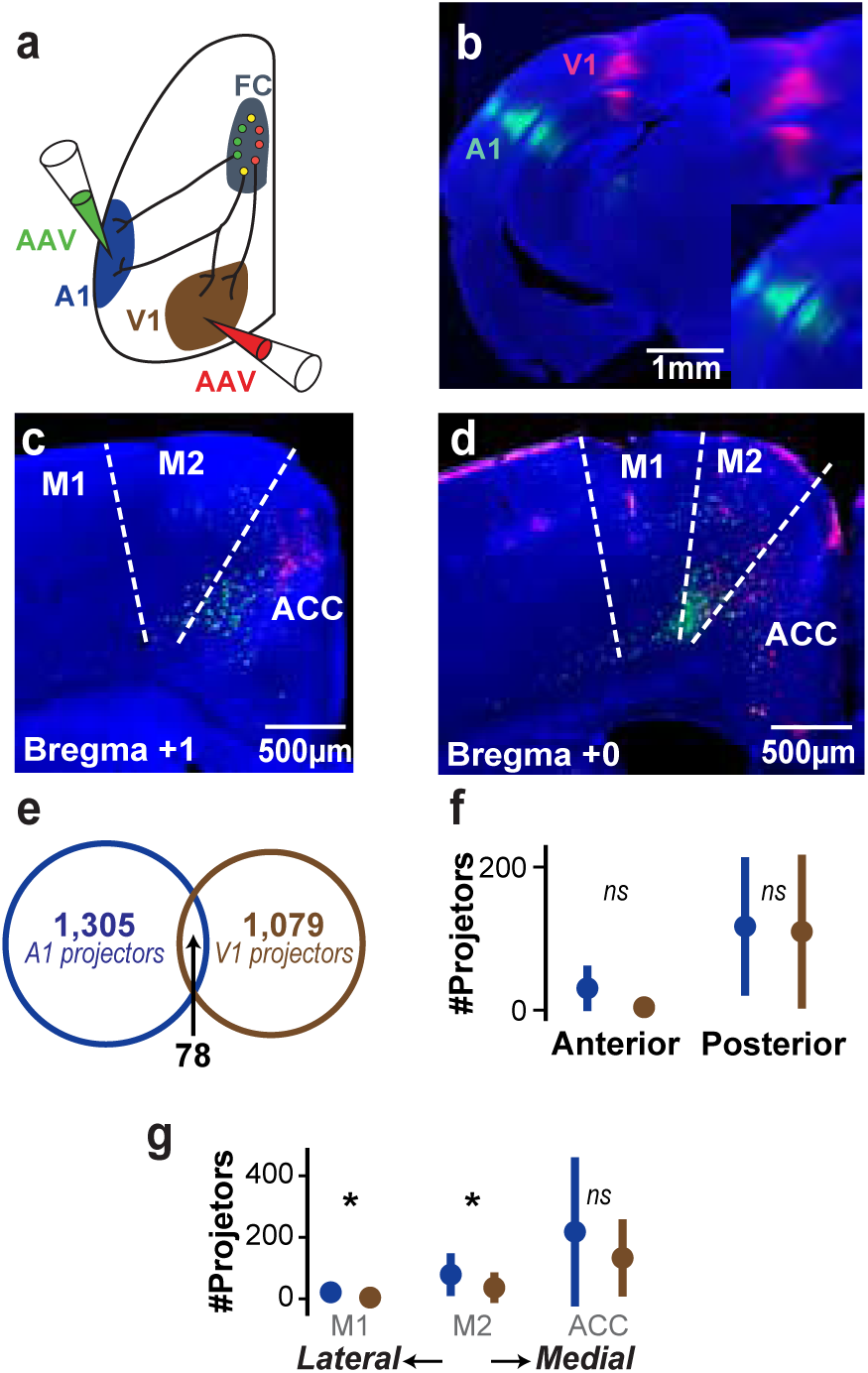
Frontal cortical neurons that project to A1 and V1 are distinct but spatially uniform. **(a)** Dual retrograde labeling of frontal cortex neurons that project to A1 and V1. **(b)** Coronal slice of the A1 (green) and V1 (red) injection sites. The insets are zoomed-in images of the same injection sites. **(c)** Example brain slice from an anterior section of frontal cortex. Cortical sub-regions were demarcated using the Paxinos atlas. **(d)** Same as ***(c)*** but for a posterior slice. **(e)** The number of frontal cortical neurons that project to A1, V1, and both (N = 3 mice, n = 54 slices). **(f)** Mean +/-sd number of frontal projectors to A1 and V1, respectively, per slice that are in anterior versus posterior frontal cortex. Statistical comparisons are pairwise, using the Wilcoxon signed rank test with a Bonferroni correction for multiple comparisons (p = 0.078 for anterior slices, p = 0.85 for posterior slices). (g) Mean +/-sd number of frontal projectors to A1 and V1, respectively, per slice along the mediolateral extent of frontal cortex. Statistical comparisons are pairwise, using the Wilcoxon signed rank test with a Bonferroni correction for multiple comparisons (M1: p = 6.13e-7, M2: p = 0.003, ACC: p = 0.074).

Given the distinct spatial organization of input from A1 and V1 to the frontal cortex, we next examined the spatial organization of the frontal projectors to A1 and V1. Generally, we observed many more sensory projectors in posterior frontal cortex compared to anterior frontal cortex, but this anteroposterior bias was consistent for both frontal projectors to A1 and V1 (Figure 6f). Along the mediolateral axis of the frontal cortex, there were more A1-projectors than V1-projectors. But generally, both projector sub-populations were biased towards the medial-extent of the frontal cortex (Figure 6g). These data thus provide evidence for a spatial organization of frontal input to sensory cortices that is modality-invariant.

### Functional organization of frontal neurons projecting to distinct sensory cortices

The anatomical segregation of the projections from the frontal cortex to A1 and V1 suggests that these pathways may also be functionally specialized. We next combined retrograde labeling with two-photon calcium imaging of frontal cortical neurons to assess the extent to which frontal cortical projectors to each sensory region encode sensory stimuli. Specifically, we injected AAVretro.Tdtomato into either A1 or V1 in separate mice to label frontal cortical neurons projecting to sensory cortex and measured their responses during auditory and visual stimulation using calcium imaging (Figures 7a-b). While we identified a few A1-projecting frontal cortex neurons responsive to sounds (Figure 7c) and V1-projecting neurons responsive to visual cues (Figure 7e), projection neurons to either A1 or V1 were largely non-responsive to sensory stimuli (Figures 7d,f). When we quantified across the population of projection neurons and sensory-responsive neurons, we found that A1-projection neurons were no more likely to be sensory-responsive than non-projection frontal cortex neurons (Figure 7g-h; sound-responsive: FC→A1 neurons = 3.7%, non-projection neurons = 6.1%, p = 0.817; visually-responsive: FC→A1 = 14.8%, non-projection neurons = 8.1%, p = 0.171, Fisher’s exact test). Similarly, V1-projecting frontal cortical neurons were no more likely to respond to either sounds or visual cues than the rest of the population (Figure 7i-j; sound-responsive: FC→V1 = 4.4%, non-projection neurons = 8.3%, p = 682; visually-responsive: FC→V1 = 8.7%, non-projection neurons = 8.0%, p = 0.560, Fisher’s exact test). Taken together, these data provide evidence for a model in which sensory information is richly encoded in frontal cortex populations, but these rich sensory signals are not preferentially conveyed back to the sensory cortex.

**Figure 7.**
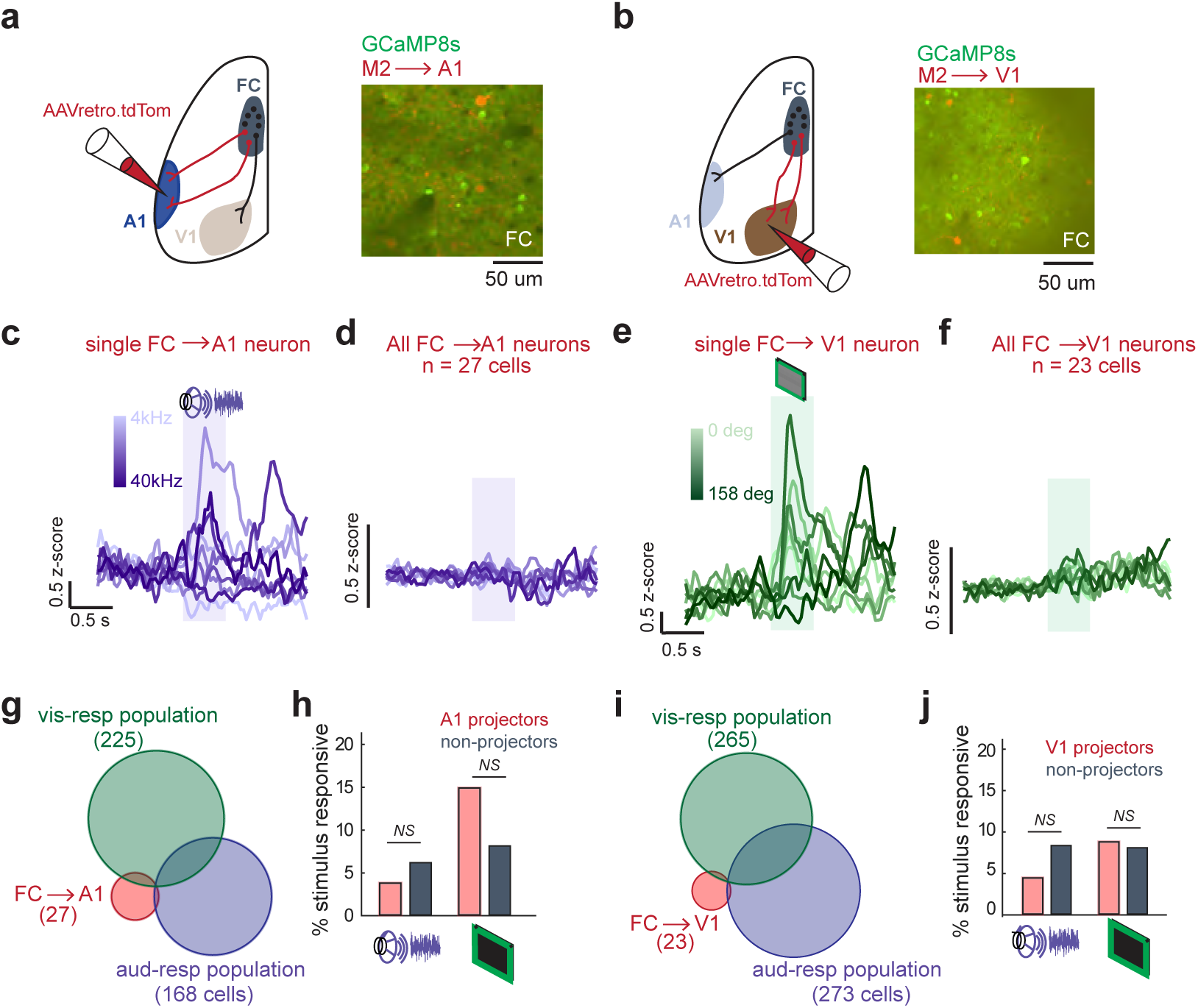
Frontal cortex neurons projecting to sensory cortex are not sensory responsive. **(a)** Left, schematic for imaging frontal cortex (FC) neurons projecting to primary auditory cortex (A1). AAVretro.CAG.TdTomato was injected into A1, guided by intrinsic imaging; right, example field of view in frontal cortex showing GCaMP8s-expressing neurons and A1-projecting FC cells. **(b)** Same as (a), but AAVretro.CAG.TdTomato was injected into primary visual cortex (V1), guided by stereotaxic coordinates. **(c)** Example sound-responsive A1-projecting FC neuron to pure tones. **(d)** Average response to each pure tone frequency of all A1-projecting FC neurons (n = 27 cells, 2 mice). **(e)** Example grating-responsive V1-projecting FC neuron. **(f)** Average response to each visual grating across all V1-projecting FC neurons (n = 23 cells, 2 mice). **(g)** Overlap between auditory-responsive, visually-responsive and A1-projecting FC cells across animals. **(h)** A1-projecting cells are no more likely to be sensory-responsive than the rest of the FC neuron population (aud-responsive: 3.7% A1 projectors, 6.1% non-projectors, p = 0.817; vis-responsive: 14.8% A1 projectors, 8.1% non-projectors, p = 0.171, Fisher’s exact test). **(i-j)** same as (g-h), but for V1-projecting FC neurons (aud-responsive: 4.4% projectors, 8.3% non-projectors, p = 0.682; vis-responsive: 8.7% projectors, 8.0% non-projectors, p = 0.560, Fisher’s exact test).

## Discussion

Although sensory processing in primary sensory cortices is known to be strongly influenced by behavioral state, movement, expectation, and other nonsensory variables via inputs from the frontal cortex, comparatively less attention has been given to the extent to which sensory information itself is represented within frontal cortical circuits. This asymmetry has contributed to a largely compartmentalized view of sensory–frontal communication, in which the encoding of sensory inputs is restricted to sensory cortex, and frontal cortex modulates sensory cortex based on behavioral and contextual variables alone. In this study, we find robust, modality-and feature-specific sensory responses in the frontal cortex that could not be fully explained by stimulus-evoked movement. Sensory inputs and responses exhibited broad but spatially structured distributions, while neurons projecting to primary sensory cortices formed largely distinct populations that were not preferentially responsive to passive sensory stimulation. By examining reciprocal frontal-sensory interactions across anatomical, population, single-neuron and spatial scales, this work addresses a fundamental gap in our understanding of how distinct sensory modalities are represented and organized within the frontal cortex. Our findings support the notion that sensory information is an intrinsic component of frontal cortical processing, where it can be integrated with behavioral and internal-state signals to support flexible behavior.

### Multimodal sensory representations in frontal cortex

Our findings reveal an underappreciated capacity of the frontal cortex to encode detailed information about the multimodal sensory environment. This extends a growing body of work showing that rodent frontal cortical regions such as M2 are not exclusively motor structures, but participate in the integration of sensory, motor, and task-related information (Murakami et al., 2014; Siniscalchi et al., 2016; Barthas and Kwan, 2017; Leinweber et al., 2017; Holey and Schneider, 2024; Zhou and Schneider, 2024; Lien and Haider, 2025; Tsukano et al., 2026; Zempolich et al., 2026). More broadly, large-scale recordings have demonstrated that sensory events are represented across distributed cortical networks, including regions traditionally associated with decision-making and action (Siegel et al., 2015; Goard et al., 2016; Musall et al., 2019; Steinmetz et al., 2019; Stringer et al., 2019). Our results add to this framework by showing that frontal sensory representations can retain information about specific stimulus features even during passive sensory presentation. Furthermore, our experiments present stimuli across different modalities and reveal that auditory and visual information is separable at the population level in the frontal cortex while a subset of individual neurons respond to both modalities. This could provide the scaffolding for the frontal cortex to integrate multimodal sensory information while maintaining modality-specific representations.

Importantly, we found that the frontal representations do not simply recapitulate those observed in primary sensory cortices. Although frontal neurons exhibited robust and feature-selective sensory responses, their response magnitudes and temporal properties differed from those measured in A1 and V1, with greater heterogeneity across the frontal population. Such differences are consistent with the idea that sensory signals are transformed as they enter higher-order cortical networks, rather than being transmitted as faithful copies of primary sensory representations (Felleman and Van Essen, 1991; Rauschecker and Scott, 2009; DiCarlo et al., 2012, 2012; Bizley and Cohen, 2013).

### Mixed coding of sensation and movement

Previous work has shown that auditory and visual responses in the frontal cortex are reduced when the auditory or visual cortex are suppressed, suggesting that stimulus-evoked activity in frontal cortex arises, in part, from direct input from primary regions (Holey and Schneider, 2024; Lien and Haider, 2025). Nonetheless, interpreting sensory-evoked activity throughout the entire brain, and particularly so for auditory-evoked activity, requires distinguishing sensory representations from the movements and changes in behavioral state that sensory stimuli themselves can evoke, especially for auditory stimuli (Bimbard et al., 2023; Clayton et al., 2024). Growing evidence that movement accounts for a substantial fraction of cortical activity, including activity in regions not traditionally considered motor areas (Musall et al., 2019; Steinmetz et al., 2019; Stringer et al., 2019) raises the possibility that the apparent sensory response in the frontal cortex could similarly reflect motor responses to the stimuli. Our GLM analysis argues against this explanation as a complete account of frontal sensory responses. Models that incorporated stimulus information in addition to measured movement better accounted for the activity of a subset of frontal neurons than models based on movement alone. Importantly, this analysis does not imply that movement and sensory representations occupy independent neural populations. Rather, it indicates that the sensory stimulus provides explanatory information about neuronal activity beyond that contained in the measured movement signal. Frontal responses therefore appear to reflect combinations of sensory and behavioral variables, consistent with a distributed view of cortical coding in which individual neurons multiplex information about external events and ongoing behavior (Rigotti et al., 2013; Fusi et al., 2016; Steinmetz et al., 2019; Stringer et al., 2019).

At the same time, our analysis places an important boundary on this interpretation. Because movement is multidimensional and cannot be captured exhaustively by a single behavioral measure, we cannot exclude contributions from unmeasured stimulus-evoked behaviors or internal-state changes. Nonetheless, the observed modality and feature selectivity of these neurons supports the interpretation that frontal cortical activity contains sensory information that extends beyond a nonspecific motor response to stimulus presentation.

### Broadly distributed but spatially biased sensory organization motif

We found that both A1 and V1 terminal fields as well as auditory-and visual-responsive neurons in the frontal cortex were broadly distributed and locally intermingled within the frontal cortex, while nonetheless exhibiting larger-scale spatial biases. Generally, this motif is commonplace in cortical circuits, such as the local salt-and-pepper organization of tone-responsive neurons despite a broad tonotopic organization within A1 (Stiebler et al., 1997; Issa et al., 2014; Kline et al., 2021). This motif is also consistent with mesoscale anatomical studies showing that corticocortical projections in the mouse are broadly distributed but spatially structured, with individual cortical areas exhibiting characteristic patterns of long-range connectivity (Oh et al., 2014; Zingg et al., 2014).

The coexistence of local intermixing and large-scale spatial bias may provide a useful organizing principle for frontal sensory processing. Local convergence of auditory and visual information could facilitate interactions across modalities, while broader spatial biases could preserve differential access to modality-specific information across frontal subregions. Multisensory processing is generally thought to depend on circuits in which information from different modalities can converge while retaining sufficient modality-specific structure to support selective behavioral weighting (Ghazanfar and Schroeder, 2006; Stein et al., 2014). Together, these results suggest that sensory information is not discarded as signals reach the frontal cortex, nor is it simply reproduced in the format established by primary sensory regions. Instead, sensory detail is retained within a broader, more heterogeneous and spatially distributed representation. Although our experiments do not directly test multisensory integration, the organization observed here provides an anatomical and functional substrate that could support both access to specific sensory features and integration across sensory streams.

### Organization of outputs from frontal cortex

We found that frontal neurons projecting to A1 and V1 were largely non-overlapping populations, providing anatomically distinct channels for communication with the two sensory cortices. In contrast to the broad but spatially biased organization of sensory inputs to the frontal cortex, these projection populations were concentrated in more posterior frontal regions, with relatively little representation in anterior frontal cortex. Moreover, the spatial distributions of A1-and V1-projecting neurons largely mirrored one another, rather than exhibiting the modality-specific biases observed for the A1 and V1 inputs. Thus, the organization of frontal-sensory communication is not a simple reflection of the sensory input pathways to the frontal cortex, but instead reflects distinct spatial architectures for incoming and outgoing information.

The lack of a direct spatial correspondence between incoming and outgoing pathways may provide an opportunity for local processing within the frontal cortex. Rather than simply relaying sensory information back to the cortical region from which it originated, sensory inputs could be distributed across a broader frontal population, where they are integrated with other sensory, motor, associative or behavioral signals before being selectively routed back to individual sensory cortices. Such an organization could allow a larger and more diverse population of frontal neurons to participate in integrating sensory information, while the resulting signals are subsequently conveyed through more restricted, modality-specific output pathways. Such an arrangement is consistent with previous studies showing that frontal feedback can influence sensory cortex according to the specific behavioral contexts (Zhang et al., 2014; Leinweber et al., 2017; Schneider et al., 2018; Holey and Schneider, 2024).

Consistent with this proposal, A1-and V1-projecting neurons were no more likely to respond to sensory stimulation than the broader frontal population. If frontal-sensory communication simply relayed sensory information back to its source, projection-defined neurons might be expected to exhibit enhanced sensory responsiveness relative to the broader frontal population. Instead, these neurons may carry signals that emerge after sensory information has been integrated with other motor, internal-state, and associative information within local frontal circuits in a context-specific manner. It would be important for future work to unveil how sensory information is transformed between its entry into the frontal cortex and the information content of the subsequent modulation of the sensory cortex.

### Sensory representations as a substrate for sensory-motor interactions

More broadly, our findings support an integrative role for selective frontal-sensory routing. Such a perspective expands the traditional view of corollary discharge, in which motor-related signals are transmitted to sensory areas to anticipate or suppress the expected sensory consequences of movement. Rather than requiring a uniform representation of movement to be broadcast across sensory cortices, frontal circuits could combine motor signals with sensory-specific information and route the resulting signals through partially distinct frontal-sensory pathways. Such an architecture could be particularly useful when a single action has modality-or feature-specific sensory consequences, as is supported by previous studies (Leinweber et al., 2017; Schneider et al., 2018; Audette et al., 2022; Audette and Schneider, 2023; Holey and Schneider, 2024; Zhou and Schneider, 2024). The sensory stimuli in these experiments were not associated with a specific movement or behavioral outcome. As such, the projection-defined neurons would not necessarily be expected to exhibit sensory responses under these conditions. However, the linking of a particular movement to an auditory or visual consequence may recruit sensory responsiveness within these same projection-defined populations, providing a potential mechanism for behavior-dependent sensory-motor integration. Determining whether sensory responsiveness emerges in these neurons when sensory stimuli become predictably coupled to specific actions will be important for distinguishing between these possibilities.

Together, these findings suggest that the sensory cortex is embedded within a broader cortical network in which sensory information is distributed to frontal circuits, integrated with other behaviorally-relevant sources of information, and incorporated into pathways that selectively communicate with individual sensory areas. This organization provides a potential circuit framework for higher-order cortical regions to integrate multimodal sensory and behavioral information while differentially influencing distinct sensory systems.

## Methods

### Experimental model and subject details

All experimental protocols were approved by New York University’s Animal Use and Welfare Committee. All experiments consisted of wild-type (C57BL/6) mice that were purchased from Jackson Laboratories and subsequently housed and bred in an onsite vivarium and kept on a reverse day-night cycle (12h day, 12h night). We used 2–4 month-old mice of both sexes (8 M and 8 F for the electrophysiological dataset, 4 M and 2 F for the histological dataset, and 2 M and 3 F for the 2-photon calcium imaging dataset) for our experiments.

### Headpost implantation

All mice were inducted under 3.5% isoflurane anesthesia vaporized in O2 and maintained at 1-1.5% isoflurane vaporized in O2 on a stereotaxic frame. Ophthalmic ointment (Optixcare) was applied to their eyes to prevent drying and their body temperature was maintained at ∼37C. 1 mg/kg Bupivacaine (Sigma-Aldrich B5274-1G) was injected around the incision site before removing the skin above the skull surface. The animal’s head was leveled in all three planes before a custom-made titanium headpost was secured to the skull using dental cement (C&B Metabond). Frontal cortex was marked at coordinates ranging from 0.3-0.7 mm anterior and 0.8-1.5 mm lateral from bregma, with experiments targeting M2 marking 0.7 mm anterior and 1.5 mm lateral to bregma and experiments targeting A24b/ACC marking 0.3 mm lateral to bregma (Leinweber et al., 2017). V1 was marked at 0.5 mm anterior and 2-2.5 mm lateral from lambda (Keller et al., 2012; Speed et al., 2020). A1 was identified via intrinsic optical signal imaging (see below). The skull was then covered with a silicone elastomer (Smooth-On) and the mice received 5 mg/kg Meloxicam injected subcutaneously for three days while recovering before any subsequent experimentation.

### Intrinsic optical signal imaging

The mice were anesthetized with isoflurane (1-1.5%) vaporized in oxygen (0.8 L/min) and kept on a heating pad set to 37°C. Muscle overlaying the left auditory cortex was excised. Mice were injected subcutaneously with acepromazine (1.5 mg/kg) approximately ten minutes prior to imaging so that isofluorane could be reduced (0.8-1%) so as to not interfere with the signal. Intrinsic signal images were acquired using a custom tandem lens microscope (composed of Nikkor 35 mm 1:1.4 and 105 1:2 lenses) and a camera (Teledyne Lumenera). Imaging was performed through a lightly thinned skull which was made transparent by saturation with 1X phosphate-buffered saline (PBS). Images of surface vasculature were acquired using green LED illumination (530 nm) and intrinsic signals were collected using red illumination (625 nm) at 4 Hz. Each trial consisted of 1 s baseline, followed by 1 s sound stimuli and 20 s intertrial interval. Images during the response period (1-2 s from sound onset) were averaged and then divided by the average image during the baseline. Image acquisition and analyses were run on custom MATLAB code. Individual auditory fields (A1, A2, AAF, VAF) were identified based on characteristic tonotopic organization determined by their responses to pure tones (1s, 75 dB; 3, 10 and 30 kHz). Injections and recordings in A1 were guided by matching vasculature patterns overlaid with the intrinsic signals (Bakin et al., 1996; Narayanan et al., 2023).

### Sensory stimulation

All sensory stimuli were controlled via custom MATLAB code using Bpod (Sanworks), leveraging PsychToolbox. Visual and sound trials were presented in random order. Sound stimuli were delivered from the computer to a sound card (RME Fireface UCX), the output of which was routed to a free-field electrostatic speaker (Tucker Davis Technologies) located ∼10 cm lateral to the mouse’s right ear. Speakers were calibrated over a range of 4-40 kHz to give a flat response (+/-1 dB). Pure tones of 8 frequencies (log-spaced, 4-40 kHz) were calculated in MATLAB at a sample rate of 192 kHZ and were presented at 70 dB SPL. Broadband white noise stimuli were generated in MATLAB and presented at 75 dB SPL. Dynamic random chord (DRC) stimuli consisted of sequences of brief tone chords spanning 4–40 kHz. Twenty-four frequencies were logarithmically spaced across this range. Individual tone pips were 20 ms in duration, including 5-ms cosine onset and offset ramps, and successive chords were presented without an intervening gap. For each chord, each of the 24 frequency components was independently included with a probability of 1/12, resulting in an average of approximately two tone components per chord. The sound level of each included component was independently selected at random from seven nominal levels ranging from 35 to 65 dB in 5-dB increments. The selected tone pips for each chord were summed to generate the chord (Linden et al., 2003).

Visual stimuli were presented using a 7 in screen (ROADOM 7’’ Raspberry Pi Screen, IPS1024×600) positioned ∼7 cm away from the mouse’s right eye, spanning ∼90 deg horizontally and ∼60 deg vertically of the animal’s visual field. In the absence of a stimulus, the screen was gray and the onset of visual stimuli was determined via a photodiode (FDS1010-CAL) positioned in the bottom right corner of the screen. Drifting gratings were at full contrast and were of one of eight orientations evenly spaced from 0-320 degrees, moving sinusoidally with a spatial frequency of 0.3 cycles/deg, and temporal frequency of 4 cycles/sec. Gratings matched the contrast of the gray screen that was shown during the inter-trial interval. Dense visual checkerboard stimuli consisted of a 20 × 30 grid of square elements that occupied the entire screen and corresponded to ∼4 deg. On each frame, each checkerboard square was independently assigned a color between white and black, generating a spatially and temporally varying visual pattern across the display. Checkerboard frames were updated at either ∼30 or ∼60 Hz (Reid et al., 1997).

All stimuli across both modalities were randomly interleaved during stimulus presentation. The inter-trial interval during the electrophysiological recordings was 2.5 seconds whereas that for the 2-photon calcium imaging recordings was 5 seconds.

### Videography and motion energy analysis

The movement of the animal during stimulus presentation in every recording was recorded using a FLIR IIS Blackfly S USB 3.1 camera (Teledyne) under constant infrared (IR) illumination. Frames were acquired at ∼66 frames/second for the duration of the recording. An additional IR LED was positioned in the background of the view of the camera. This LED was triggered to flash at the start of each trial via the BPOD program, enabling alignment of the videography data to the electrophysiology data. The motion energy of a region of interest (ROI) caudal to the vibrissa array was manually identified (Clayton et al., 2024) and a separate ROI outlining the triggering IR LED was extracted using facemap (Syeda et al., 2022). Custom MATLAB scripts then extracted the facial motion energy of each trial from the first ROI, identified by the flashes of the triggering IR LED measured in the motion energy of the second ROI. The motion energy of each trial was quantified by computing the area under the curve during each 500 ms stimulus presentation.

### Electrophysiological recordings

All mice were habituated to head-fixation for a minimum of 2 days, with the duration of the head-fixation gradually increasing each day (Guo et al., 2014). Once habituated, a small craniotomy was made to expose the brain surface over the marked brain region of interest. Briefly, all mice were inducted under 3.5% isoflurane anesthesia vaporized in O2 and subsequently maintained at ∼1.2% isoflurane vaporized in O2 on a stereotaxic frame. Ophthalmic ointment (Optixcare) was applied to their eyes to prevent drying and their body temperature was maintained at ∼37C. The skull was then thinned above the region of interest before exposing the brain while leaving the dura intact. The exposed brain was then covered with a silicone elastomer (Kwik-Sil, WPI), a silicone elastomer, and given a minimum of 2 hours to recover.

The mice were then head-restrained on a custom platform made of ThorLabs parts, placed in the dark in a sound attenuating booth (Gretch-Ken). The Kwik-Sil was removed and the brain surface kept moist with saline. A 128-channel probe (Neuronexus, A3×43_42_43-edge) was inserted to the depth of interest using an electronic manipulator (Sensapex) at a speed of 2 µm/sec. Recordings targeting ACC were lowered to ∼2 mm while all other cortical recordings were lowered to a depth of 0.9-1 mm. Upon reaching the targeted recording depth, we waited a minimum of 10 min for the brain to settle before starting the recording and presenting a minimum of 20 trials of each stimulus, randomly interleaved across stimuli. Only one recording was performed per day and up to three recordings were performed per craniotomy on subsequent days, after which an additional craniotomy was made under the same conditions to target an additional brain region of interest.

The 128-channel recording probe was connected to 2 64-channel digitizing headstages (Intan). All continuous signals, including those from the recording probe, digital triggers from BPOD to indicate the start of each trial, and the analog signal from the photodiode were acquired at 30 kHz, monitored in real-time, and saved for offline analysis using OpenEphys software. Each stimulus type was presented 20 times and all stimulus types were randomly interleaved.

### Electrophysiological data analysis

The action potentials were extracted and assigned to individual neuron clusters using Kilosort4 for automated spike sorting based on template-matching (Pachitariu et al., 2023, 2024)a nd manually curated using Phy2 (Rossant and Harris, 2013). A custom MATLAB script then processed the curated clusters to compute the various metrics to assess the quality of each neuron cluster (Dimwamwa et al., 2024). Only well-isolated, single neuron clusters were used for subsequent analysis, identified by the following: a signal-to-noise ratio of the mean spike waveform greater than three and less than 2% of all spikes violating a 2 ms imposed refractory period, and a coefficient of variation of the spiking over the duration of the recording less than 1 (using 120 second long bins).

The sensory response of each neuron was quantified by measuring the average baseline-subtracted firing rate across trials, using spikes occurring within 500 ms of the stimulus subtracted by the pre-stimulus spiking occurring in a window of equivalent size on a trial-by-trial basis. To identify stimulus-responsive neurons, we compared the post-stimulus spike counts in the 500 ms window following the relevant stimulus to an estimate of the neuron’s spontaneous spiking using a permutation test on Cohen’s *d*. The spontaneous activity was estimated using the pre-stimulus spike counts of all stimuli in an equivalent window. We generated a null distribution by pooling and randomly reassigning the label of the spontaneous and evoked spike counts 10,000 times, and computing the Cohen’s d each time. Then a two-sided permutation p-value was calculated from the proportion of shuffled Cohen’s d values with an absolute magnitude greater than or equal to the observed value. Neurons were considered responsive at p < 0.01.

For neurons identified as stimulus-responsive, we computed the fano factor as the ratio of the variance of the baseline-subtracted spike counts across trials to the mean. For neurons identified as stimulus-responsive and responding with an increase in spiking, the onset latency was computed from the cumulative spike count using a bootstrap procedure to compare to spontaneous spiking. Stimulus-evoked and trial-averaged spike trains (beginning at stimulus onset) were binned at 1-ms resolution and summed cumulatively for the duration of the trial. Spontaneous activity was estimated by drawing with replacement pre-stimulus spike trains for 1,000 bootstrap iterations. The onset latency was defined as the first post-stimulus time bin at which the observed cumulative evoked response exceeded the 99% upper confidence bound of the spontaneous distribution (Wiest et al., 2005; Pala and Stanley, 2022).

Population-level dynamics were assessed using principal component analysis. For each neuron, PSTH responses were z-scored across all time bins and stimulus conditions by subtracting the neuron’s mean activity and dividing by its standard deviation. The normalized PSTHs were concatenated across stimulus conditions to generate a #condition*time-by-#neurons matrix. PCA was performed using MATLAB’s *pca* command after mean-centering each neuronal dimension, and the population activity at each time point and stimulus condition was projected onto the resulting principal component axes. The percentage of total population variance explained by each principal component was quantified from the corresponding PCA eigenvalues.

The auditory noise from visual flash stimuli were decoded from populations of individual neurons using a linear support vector machine (SVM) applied on the trial-by-trial, baseline-subtracted spike counts in a 500 ms window. Decoding was performed using MATLAB’s *fitcsvm* command on population sizes ranging from 5 to 200 neurons. For each population size, neurons were randomly sampled from the full recorded population, and 25 independently sampled neuronal populations were used to estimate the average classification performance. Classification was performed using a stratified five-fold cross-validation scheme with equivalent numbers of auditory and visual trials in each fold. Feature normalization was performed independently within each cross-validation fold using the mean and standard deviation of the training data, and the resulting parameters were then applied to the held-out test data. Statistical significance was assessed using a permutation test in which stimulus labels were randomly shuffled across trials and the complete decoding procedure was repeated 1,000 times to generate a null distribution of decoding accuracies. The permutation *p*-value was calculated as the proportion of shuffled decoding accuracies greater than or equal to the observed decoding accuracy.

To determine whether neural population activity encoded the identity of individual auditory and visual stimuli, we performed multiclass decoding of the eight auditory tones and eight visual gratings using an SVM using MATLAB’s *fitcecoc* command. For each trial, baseline-subtracted spike counts measured within a 500-ms stimulus-response window were used as normalized features as described above, with each stimulus assigned a separate class label. Again, decoding performance was evaluated using stratified five-fold cross-validation, such that each fold contained an equivalent number of trials from each stimulus class. Classification accuracy was calculated as the proportion of correctly identified held-out trials, with chance performance equal to 1/16 (6.25%).

### Generalized Linear Model (GLM)

The spiking activity of each neuron, stimulus timing for each trial, and the associated motion energy were organized in Neurodata Without Borders (NWB; (Rübel et al., 2022) format using custom MATLAB scripts. Generalized linear models (GLMs) were then fit separately to the spiking activity of each neuron using NeMoS, a statistical modeling framework optimized for systems neuroscience (Balzani et al., 2025). The structure of each GLM was a Poisson observation model with a log link function (Nelder and Wedderburn, 1972; Pillow et al., 2005). For each sensory modality, two model types were fit: a movement-only model containing motion energy as the sole predictor and a full model containing both movement and stimulus predictors. Models were fit using 40 trials comprising 20 catch trials during which no stimuli were presented and 20 stimulus trials. Auditory and visual trials were modeled separately; therefore, each neuron was fit with six models in total: three for the auditory comparison and three for the visual comparison. All models were fitted using a five-fold cross validation scheme where each fold contained an equal number of catch and stimulus trials. We evaluated the performance of each model using McFadden’s pseudo-R^2^ score applied to the log-likelihood of each model. Ultimately, only sensory responsive neurons with model scores greater than 0 were included for analysis.

Each trial used for model fitting was restricted to a window beginning 1 s before stimulus onset and extending through the 500-ms stimulus-presentation period, thereby excluding stimulus-offset responses. Spiking activity and motion energy were discretized into 5-ms bins and the motion energy was z-scored. The stimulus was represented by two binary predictors: an impulse marking stimulus onset and a boxcar predictor spanning the duration of stimulus presentation to capture the sustained response. The stimulus and movement predictors were convolved with five raised-cosine basis functions with a 250 ms history (Pillow et al., 2005). The models also incorporated ridge regularization. Regularizer strength, type of basis function, and number of basis functions were selected by grid search using only the training data of each cross-validation fold.

### Anatomical tracing data acquisition

After headpost implantation and intrinsic optical signal imaging (see above), small craniotomies were made above A1 and V1. Micropipettes that were 17.5-22.5 µm in diameter were pulled using a pipet puller (P-97, Sutter Instruments) and loaded on the microinjector (Nanoject). Pipettes were lowered to 875 µm at 2 µm/s, given 5 minutes for the brain to settle, and then retracted to 800, 500, and 270 µm for each injection. At each depth, 5nL of virus was dispensed per cycle at a rate of 5 nL/s, with 30 second intervals between each cycle. For injections into A1, the microinjector was positioned 30° from vertical in the mediolateral plane. To maximize spread within A1, 40 nL of virus was injected at each depth in two separate craniotomes that were 150-200 um apart (120 nL per craniotomy, 240 nL total). For injections into V1, the microinjector was positioned perpendicular to the skull surface and 80 nL of virus was injected per depth (240 nL total).

For anterograde tracing experiments, mice received injections of rAAV1-CAG-GFP (4.7E12; Addgene) and pAAV9-hSyn-mCherry (2.4E12; Addgene) into A1 and V1, respectively, with fluorophore identity counterbalanced across animals. Similarly, for retrograde tracing experiments, mice were injected with retro-pAAV-hSyn-EGFP (19E12; Addgene) in A1 and retro-AAV-hSyn-mCherry (2.1E13; Addgene) in V1, respectively, with fluorophore identity again counterbalanced across animals. After a minimum of 21 days, the mice were then anesthetized and transcardially perfused with saline and 4% paraformaldehyde. The brain was removed and post-fixed in 4% paraformaldehyde overnight. The following day, the caudal pole was glued to the base of a Leica vibratome and cut into 70 µm slices. Slices were incubated in DAPI (1ug/mL dissolved in 1X PBS, Sigma-Aldrich D9541-1mg) for 10 minutes before being mounted onto slides using aqueous mounting medium with DAPI (Abcam ab 104139). Slides were scanned in an Olympus VS200 at 4x resolution using 358, 495, and 544 nm color channels.

### Anatomical tracing data analysis

The Anterograde tracing data was analyzed using custom MATLAB scripts. The frontal cortex regions of interest (ROIs) were identified in each slice using the Franklin and Paxinos, 3rd edition atlas (Franklin and Paxinos, 2008). The red and green color channels in the ROIs were separately thresholded and binarized before applying a radial sweep extending from the border between M2 and A24b was applied to account for the curvature of the cortex at the medial edge. The thresholded axon density for each color channel was binned into 5° sections and plotted as a function of angle away from the M2/A24b boundary.

The retrograde tracing data was analyzed using NeuroInfo (Tappan et al., 2019). We first used the software to align and transform images of slices containing the frontal cortex to fit the Allen Common Coordinate Framework (Wang et al., 2020) before performing cell counting, with manual adjustments made to remove artifacts. Custom scripts utilized the software output to plot cell counts per region along the anteroposterior and mediolateral axes.

### Two-photon calcium imaging recordings

A craniotomy (3 x 3 mm^2^) was made over the left frontal cortex, leaving the dura intact. Stereotaxic markings were made at coordinates for M2 (+1.5mm Anterior, 0.7mm Lateral from Bregma), which allowed for localization and alignment across animals. Drilling was interrupted every 1-2 seconds and the skull was cooled with phosphate-buffered saline to prevent damage from overheating. Virus was injected at 8-10 locations (250 µm deep from the pial surface, 40 nL/site at 1 nL/sec). AAV9.syn.GCaMP8s.WPRE.SV40 was injected into WT B6 mice. A glass window was placed over the craniotomy and secured with dental cement (C&B Metabond). 5 mg/kg Meloxicam was administered immediately after surgery and two additional days post-op. For retrograde labeling experiments, AAVretro.CAG.tdTomato was injected in either A1 or V1 prior to making the frontal cortex craniotomy. Two-photon calcium imaging was performed 2-3 weeks after chronic window implantation to ensure an appropriate level of GCaMP8s expression. On the day of imaging, awake mice were head-fixed under the two-photon microscope and responses to visual and auditory cues were measured. GCaMP8s was excited at 925 nm (Insight X3, Newport), and images (FOV: 450×625 µm, 512×705 pixels) were acquired with a custom two-photon laser scanning microscope (Neurolabware) running accompanying custom MATLAB software (Scanbox) using a x16 objective (Nikon) at 30 Hz. Images were acquired from L2/3 (200-250 µm below the surface). Timings of stimulus delivery were aligned to imaging frames by recording the timing of 5V trigger signals sent by the stimulus delivery computer. Stimuli were presented for 0.5 s, and total trial duration was 5 s to allow activity to return to baseline. Video of each animal’s face and body were recorded to allow for post-hoc analysis of contributions of movements to the neuronal response. Auditory and visual stimuli were presented in interleaved trials by a computer running Bpod (Sanworks). A photodiode was placed at the bottom corner of the visual stimulus delivery screen to accurately account for delays between stimulus delivery commands and screen refreshing.

### Two-photon calcium imaging data analysis

Lateral motion during imaging was corrected using Suite2P software (https://github.com/Mouseland/suite2p) (Pachitariu et al., 2017). ROIs corresponding to individual cell bodies were also automatically detected by Suite2P software and supplemented by manual curation. The true somatic fluorescence signal was estimated as dF(t) = F_measured_(t) - 0.9 × F_background_(t). Normalized dF/F time series were subsequently generated from a smoothed baseline. ROIs were judged as significantly excited if they fulfilled the following criteria: (1) unsmoothed dF/F had to exceed a fixed threshold value (2.0 × standard deviation of the baseline) consecutively for at least 3 frames in at least 50% of trials for at least one stimulus. (2) dF/F averaged across trials had to exceed the same fixed threshold value consecutively for at least 3 frames for at least one stimulus. ROIs that were not at least 3% brighter than the background were excluded from further analyses. Auditory-responsive and visual-responsive cells were determined if they met these criteria for at least one frequency or drifting grating, respectively. Cells responsive to either modality, but not both, were included for the spatial analyses used in this study. In the retrograde labeling experiments, frontal cortex cells projecting to A1 or V1 were labeled with TdTomato and matched with GCaMP8s-expressing cells using custom Matlab code.

For stitching of imaging FOVs, images were overlaid by matching vasculature and morphology and stitched with custom MATLAB code. To compare spatial distribution of sensory-responsive cells across mice, Anterior-Posterior distance and Medial-Lateral distance were normalized. To evaluate local spatial organization, the fraction of same-modality nearest neighbors was computed for each neuron across a range of neighborhood sizes (k = 1-25). Observed mixing fractions were compared against an empirical change null distribution generated via spatial label permutation (n = 500 shuffles).

## Statistical analysis

Unless otherwise stated, the statistical significance of all comparisons were evaluated using either the two-sided Wilcoxon Ranksum test for independent pairs or the two-sided Wilcoxon Signed Rank test for dependent pairs. In two-photon imaging experiments, differences in proportions between groups were assessed using Fisher’s exact test, which was used to accommodate unequal sample sizes between groups. Differences between spatial distributions of auditory-and visually-responsive cells were evaluated using the two-sample Kolmogorov-Smirnov test.

## Acknowledgments

We thank members of the Schneider lab for input and advice throughout the project. We thank the NYU IT High Performance Computing resources, services, and staff expertise. In particular, we thank Shenglong Wang for his assistance. We also thank the staff of the Simons Institute Center for Computational Neuroscience. In particular, we thank Edoardo Balzani for his assistance. This work was supported by the NIDCD (R01-DC018802, D.M.S; F32-DC022169, A.M.K). D.M.S. was supported by a McKnight Scholar Award. D.M.S. is a New York Stem Cell Robertson Investigator. E.D. was supported by the Simons Foundation Society of Fellows.

---

**Figure S1.**
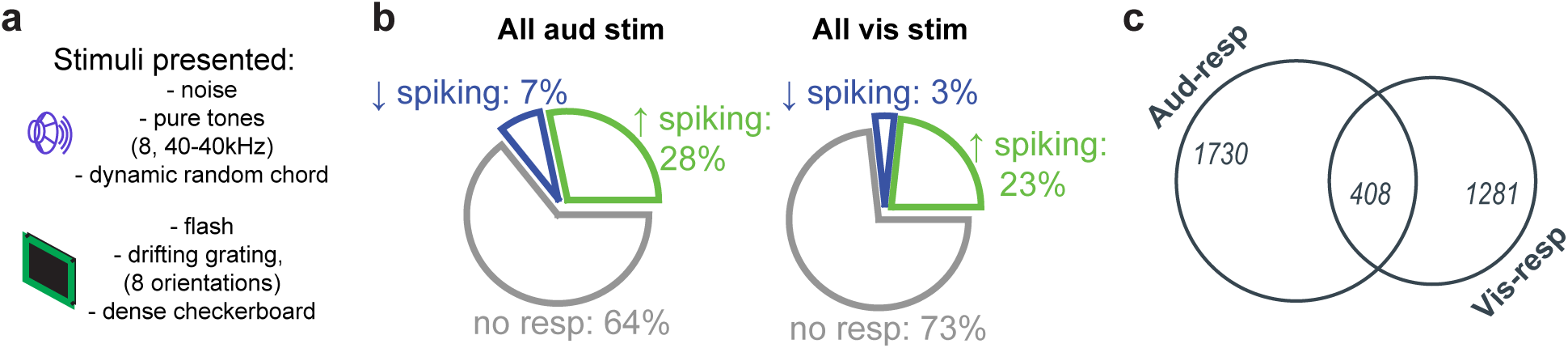
Frontal cortical neurons distinctly encode auditory and visual input across simple and complex auditory stimuli. **a.** In addition to the auditory noise stimulus and visual flash stimulus, we also presented other simple and complex stimuli. **b.** Evaluating sensory-responsiveness using the response to any of the presented simple and complex stimuli increases the proportion of responsive frontal cortical neurons compared to that presented in Figure 1. **c.** Even when factoring the response to other stimuli, a large proportion of sensory responsive frontal neurons respond to stimuli of a single modality.

